# Human Microbial Transplant Restores T Cell Cytotoxicity and Anti-Tumor Response to PD-L1 Blockade in Gnotobiotic Mice

**DOI:** 10.1101/2020.08.07.242040

**Authors:** Joshua N. Borgerding, Joan Shang, Graham J. Britton, Hélène Salmon, Camille Bigenwald, Barbara Maier, Samuel R. Rose, Ilaria Mogno, Alice O. Kamphorst, Miriam Merad, Jeremiah J. Faith

## Abstract

Recent studies demonstrate that gut microbiota regulate tumor response to immune checkpoint blockade. Still, the mechanisms by which microbiota control tumor response to immunotherapy remain unclear. We colonized germ-free mice with cultured human-derived microbiota prior to tumor inoculation. While no human donor microbiota altered tumor growth, two distinct gut microbiota inhibited tumor response to anti-PD-L1. Colonization with non-responder microbiota led to reduced tumor immune cell infiltration and modified antigen presenting cell phenotype. RNA sequencing of tumor-infiltrating CD8+ T cells revealed enrichment for stem cell-like genes in non-responders and reduced effector-like expression conferring cytotoxic potential. Antibiotic depletion and microbiota transplant restored anti-PD-L1 response in non-responders, with expansion of effector cells and cytotoxicity. Concomitant blockade of TNFα similarly improved response to anti-PD-L1 and increased cytotoxicity. These results demonstrate inhibitory roles for the microbiota in checkpoint blockade and reveal the potential for microbiota transplant and TNF blockade to overcome microbiota-mediated resistance.

## Introduction

Chronic antigen exposure generates exhausted CD8^+^ T cell responses in the immunosuppressive tumor microenvironment [1]. Antibodies directed against PD-1 and PD-L1 have shown great success in the clinic and work similarly to restore cytotoxic function to exhausted CD8^+^ T cells [2] through *cis*-regulation of the costimulatory molecule CD28 [3, 4]. Single cell RNA sequencing and ATAC sequencing have demonstrated two major subsets of exhausted CD8^+^ T cells within the tumor microenvironment [5]. The progenitor-like (or stem-like) exhausted CD8^+^ cells are polyfunctional, producing cytokines like IL-2 and TNFα, maintaining both the ability to self-renew and differentiate into effector T cells. The exhausted terminal effector CD8^+^ subset has more limited cell fate and enhanced cytotoxic potential [6]. Because they can maintain the T cell pool, progenitor-like exhausted cells are necessary for a durable anti-PD-1 effect on the tumor [7].

Immune checkpoint blockade therapies have improved survival for patients with a number of solid and hematopoietic malignancies [8], but despite dramatically altering the clinical course in a subset of patients, these drugs fail to elicit durable response for the majority of cancer patients [9]. Reports further suggest that while combination therapies increase response over single agents, they are also associated with significantly more autoimmune toxicity [10]. There is therefore great need to identify tools to reverse non-response. Human intestinal microbiota are unique and stable communities that regulate immune function in various models of disease and are causally associated with several immune-mediated conditions [11]. The ability of the gut microbiota to regulate immune activity both locally at mucosal sites and systemically makes it an attractive therapeutic target. Several studies have demonstrated that intestinal microbiota can regulate response to anti-PD-1 therapies, and microbial taxa have been identified that correlate with checkpoint blockade response [12-15]. The mechanisms through which the microbiota alter immune activity to clinical effect remain unclear. Moreover, the bacterial taxa associated with PD-1 responding patients have varied between studies, likely owing to differences in study design and patient populations [16]. Regulation of host immunity by the microbiota often occurs at the strain level [17, 18], detail not captured by 16S sequencing, which has been used to find associations between gut microbiome composition and checkpoint response in human studies [12, 13, 15].

We have previously evaluated the role of human-derived microbiota on intestinal immune modulation and induction of colitis [17, 18]. Here, we screened seven culture-defined microbiota isolated from human fecal donors for their ability to influence B16 tumor response to anti-PD-L1. While prior studies focused on microbiota to enhance anti-tumor response, we identified two communities that impaired tumor response to checkpoint blockade. To find mechanisms of non-response, we studied in detail two microbiota, which confer distinct naïve immune phenotypes and tumor immune response to anti-PD-L1. We demonstrate that microbiota-induced non-response to anti-PD-L1 is associated with reduced immune infiltration, limited effector CD8^+^ T cell proliferation and differentiation, and impaired T cell killing ability combined with increased TNFα production by myeloid cells. These immune defects could be altered and tumor response restored by combination therapy of anti-PD-L1 and anti-TNFα or transplantation of responder bacteria prior to anti-PD-L1 administration.

## Results

### Human Microbiota Can Inhibit Checkpoint Blockade Response and Alter Gut and Skin Immune Tone

To evaluate the influence of the gut microbiota on B16-F1 melanoma tumor response to checkpoint blockade with anti-PD-L1 in C57Bl/6 mice, we explored a range of different gut microbiotas including SPF mice from different vendors (Jackson Labs [Jax] and Taconic Farms [Tac]), microbiota-depleted mice (germ-free or antibiotic treated), and gnotobiotic mice colonized with each of seven different human gut microbiota cultured collections (fig. S1A, Supplementary Table 1). We did not find a microbiota-dependent difference in tumor growth of untreated mice (p=0.99; ANOVA; fig. 1A).

**Figure 1.**
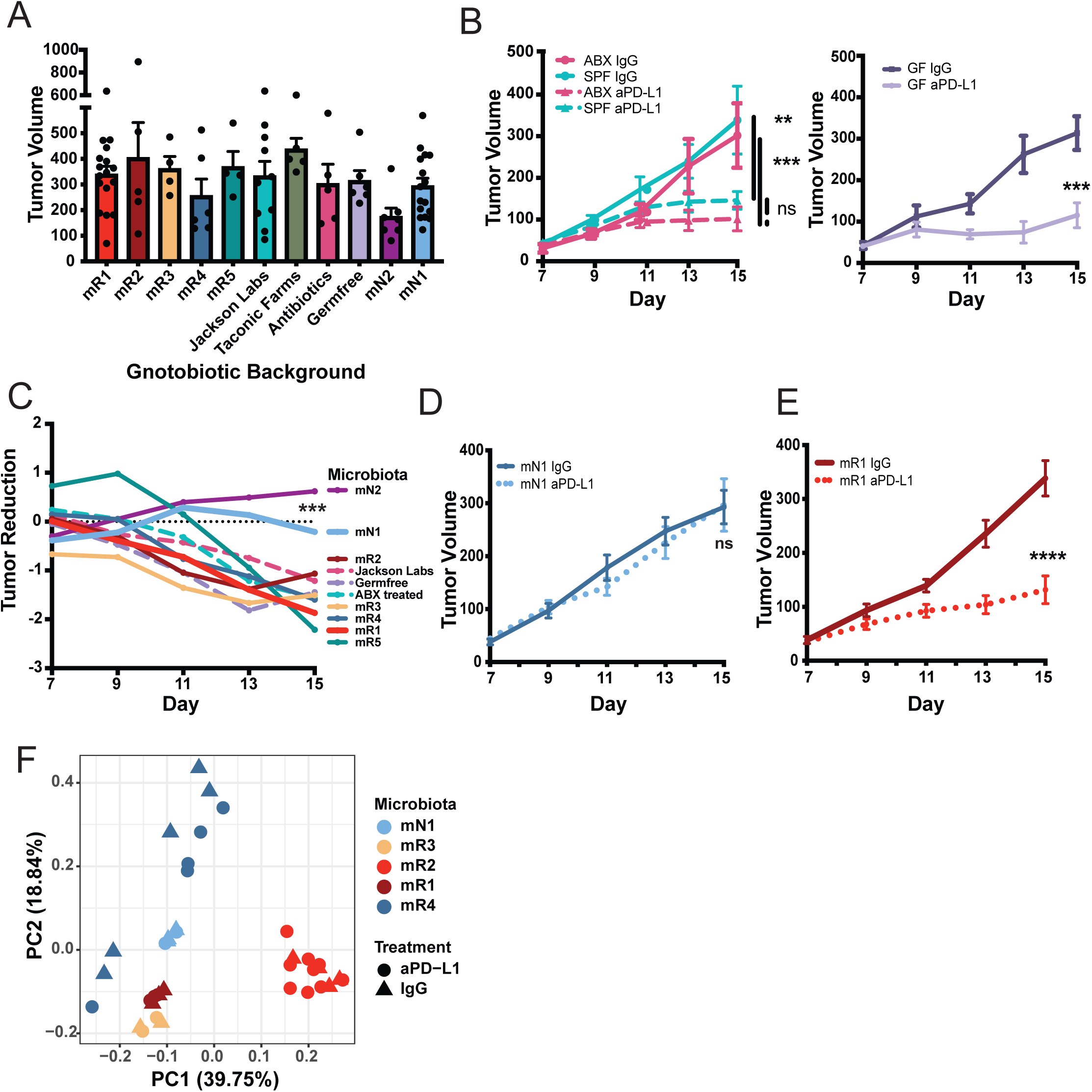
Human Intestinal Microbiota Inhibit Anti-Tumor Response of PD-L1 Blockade. (A) B16 tumor volume at day 15 in IgG isotype-treated mice, mean ± SEM. Data representative of at least two independent experiments per microbiota. p=0.99; One-way ANOVA. (B) SPF mice from Jackson Laboratories were treated with oral antibiotics for two weeks before injection of B16 tumor cells. Mice were continued on antibiotics throughout the experiment and treated with anti-PD-L1 on days 7, 9, and 11. Germfree status was confirmed by fecal culture. Data represent mean ± SEM of two independent experiments, two-way ANOVA. (C) The log_2_ median ratio in tumor volume of anti-PD-L1 treated tumors to isotype treated tumors was assessed over the course of tumor development. (D) Individual tumor growth for mN1 nonresponder mice. (E) Individual tumor growth for mR1 responder mice. P-value determined from two-way ANOVA of 4 independent experiments in head-to-head comparison with (D). (F) Fecal metagenomic analysis of gnotobiotic mice at sacrifice after either IgG or anti-PD-L1 treatment. *p<0.05, **p<0.01, ***p<0.001, ****p<0.0001

B16-F1 tumor response to anti-PD-L1 has not been evaluated in microbiota-depleted or germ-free conditions. We found that treatment with PD-L1 blockade reduced B16 tumor growth similarly in both antibiotic-treated and germ-free mice compared to SPF controls, suggesting that the microbiota was not required for anti-PD-L1 efficacy in this model (fig. 1B). This was consistent with previous results showing that anti-PD-1 antibody therapy results in significant tumor reductions in germfree mice bearing MC38 and MCA-205 tumors [13, 14], but this varies from the demonstration that anti-CTLA-4 was ineffective for treating MCA-205 sarcoma in germ-free animals [19]. Among the gnotobiotic mice colonized, two microbiota, mN1 (fig. 1C and 1D) and mN2 (fig. 1C and S1B), impaired tumor response to anti-PD-L1. Mice colonized with the other five defined microbiota (mR1-mR5) responded similarly to germ-free, antibiotic, and SPF mice (figs. 1C, 1E, S1B).

All of the gut microbial communities are comprised largely of anaerobic bacteria, with similar species overlap among pairwise comparisons of responder-responder (19.8%±8.2%), nonresponder-nonresponder (5.3%), and responder-nonresponder (12.2%±3.9%) (supplementary tables 2 and 3). The gut microbial composition of each of the seven defined gut communities was largely unchanged following anti-PD-L1 therapy (fig. 1F, S1C), although we found that mR4 had significant change in community composition (fig. S1D, p=0.043, ANOVA) in one species (*Bifidobacterium breve*; p=0.009, t-test).

To interrogate the immune factors driving differences in B16-F1 tumor response to anti-PD-L1 between gnotobiotic mice, we performed experiments focused on non-responder, mN1 (fig. 1D), and responder microbiota, mR1 (fig. 1E). In tumor-naïve gnotobiotic mice colonized with these communities, we found a disparity between the gut and skin immune phenotypes. In the colon lamina propria, the responder microbiota induced a more inflammatory profile, with higher TNFα production by antigen presenting cells and fewer regulatory T cells (fig. S1E). In the skin, however, the inflammatory relationship was reversed with the nonresponder microbiota colonized mice demonstrating more inflammatory skin immune profiles, characterized by higher TNFα production within both dendritic cell subsets and CD4^+^ T cells (fig. S1F).

### Nonresponder Community, mN1, Confers Reduced Immune Recruitment and APC Activation

To explore potential immune factors that might explain differences in tumor response to anti-PD-L1 between mN1 and mR1 mice, we performed immune profiling of the tumor and tumor draining lymph node at day 10 after tumor implantation. This enabled us to study the early development of the immune response on therapy without the confounding variable of disparate tumor size seen at the day 15 response endpoint. Tumor size has been shown to be a large driver of immune recruitment, activation, and response to therapy [20]. We found that mN1 colonized mice treated with anti-PD-L1 had reduced recruitment of CD45^+^ cells (fig. 2A), with a trend towards decreased T cells (fig. 2B), driven by CD8^+^ T cells (fig. 2C). Unlike mR1 mice, this recruitment corresponded to no change in the CD8/T_reg_ ratio with anti-PD-L1 treatment (fig. 2D). These results are supported by numerous prior studies demonstrating that increased tumor immune infiltration and increased CD8/T_reg_ ratio are predictive of response to checkpoint blockade agents [15, 21-23].

**Figure 2.**
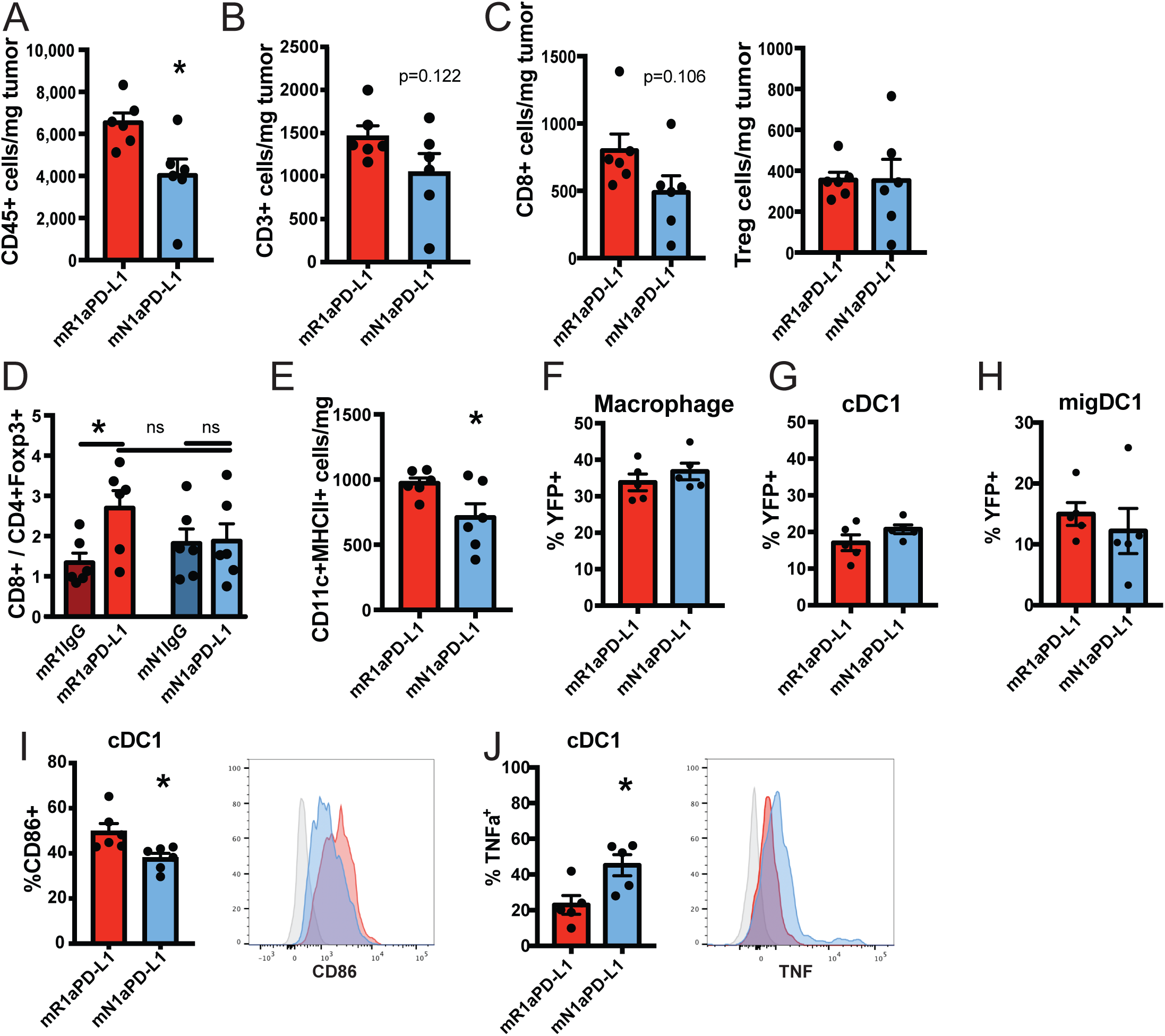
Nonresponder Microbiota Reduces Immune Infiltration and Modifies Dendritic Cell Phenotype. (A) CD45 cell count in the tumor divided by tumor mass. Data representative of two experiments, mean ± SEM. (B) CD3 T cell density at day 10 from two experiments, mean ± SEM. (C) CD8 T cell and T_reg_ density at day 10 from two experiments, mean ± SEM. (D) CD8/T_reg_ ratio. (E) Density of antigen presenting cells, mean ± SEM. (F) B16-YFP tumors were injected and the frequency of YFP positive cells in the given population was calculated by FACS for cDC1 and (G) macrophages. (H) YFP positive migratory (MHCII^hi^CD11c^int^) cDC1 in the tumor draining lymph node, mean ± SEM. (I) CD86 expression on tumor cDC1, mean ± SEM. Representative histogram shows isotype control in grey and anti-PD-L1 treated mR1 and mN1 in red and blue respectively. (I) Cells from the tumor were incubated for 4 hours in brefeldin A to measure baseline TNFα production. All data collected at day 10 during anti-PD-L1 treatment. Data represented as mean ± SEM for at least two independent experiments *p<0.05.

Antigen presenting cells (APCs) are critical components of an effective anti-tumor immune response owing to their ability to prime naïve T cells. We and others have shown that strategies to increase the number of APCs (specifically type 1 dendritic cells, cDC1), can be used to enhance effectiveness of anti-PD-L1 [21, 24]. We found mN1 colonized mice had a lower abundance of CD11c^+^MHCII^+^ cells in the tumor compared to mR1 mice (fig. 2E) with no unique difference between relative proportions of macrophages, inflammatory monocytes, or dendritic cells that comprise the CD11c^+^MHCII+ APC compartment (fig. S2B).

Within the tumor environment, antigen presentation to CD8^+^ T cells requires effective uptake of tumor antigens and delivery to the tumor-draining lymph node. While both macrophage and cDC1 can activate T cells at the tumor site, cDC1 are uniquely able to transport intact tumor antigen to the tumor draining lymph for priming of naïve T cells, necessary for effective tumor immunity [21, 25]. When we implanted B16-tumor cells expressing YFP, we found no difference in the proportion of macrophages (fig. 2F) or cDC1 (fig. 2G) in their uptake of tumor antigen, or the frequency of migratory cDC1 that delivered tumor antigen to the draining lymph node (fig. 2H). To activate T cells, APCs must express costimulatory molecules on their cell surface. In particular, antagonism of the PD-1/PD-L1 pathway relies on modification of the CD28-CD80/CD86 axis [3]. Supporting the role of cDC1 differences potentially driving the lack of response in mice colonized with the mN1 microbiota, we found cDC1 in mN1 mice expressed less CD86 relative to the mR1 colonized responder mice (fig. 2I) with no significant differences for CD86 in macrophages (fig. S2F) or CD80 expression by either cDC1 (fig. S2C) or macrophages (fig. S2F). T cell activation also requires cytokine secretion from APCs to direct activity [26, 27]. While cDC1 and macrophages from tumors of mN1 and mR1 mice had no difference in IL-1b or IL-12 production (figs. S2D, S2G), mN1 tumor APCs produced significantly more TNFα (figs 2J, S2H). In the tumor draining lymph node, there was no difference in costimulatory expression, IL-12 or IL-1b production, but migratory cDC1 again produced more TNFα in mN1 tumor-bearing mice (fig. S2E). Taken together these results demonstrate that the intestinal microbiota can regulate tumor immune infiltration, alter key APC subsets in the tumor associated with response to anti-PD-L1, and perturb APC cytokine production in both the tumor and tumor draining lymph node.

### mN1 Community Impairs Expansion of Effector CD8^+^ Cells and Reduces Cytotoxicity, Reversible by TNFα Blockade

Effective anti-PD-L1 therapy requires CD8^+^ T cells [28], with an inhibitory role for CD4 cells in mouse tumor models [29]. We depleted CD4 cells to evaluate the importance of this compartment for microbiota-induced anti-PD-L1 failure (fig. S3A). While CD4 depletion alone reduced tumor growth in mR1 mice, it also enhanced tumor control in combination with anti-PD-L1 (fig. S3B). In mN1 mice however it had no significant impact alone or in combination (fig. S3C). We therefore focused our attention on changes to the CD8^+^ T cell compartment to explain anti-PD-L1 failure in mN1 mice. To evaluate the generation of an antigen specific response, we stained for SIINFEKL^+^ T cells in B16-ova bearing mice. We found no difference in frequency of ova-specific T cells in either the tumor or tumor draining lymph node (fig. 3A). This suggested that phenotypic changes to T cells, rather than differences in antigen-specific T cell number, were responsible for anti-PD-L1 failure in mN1 mice.

**Figure 3.**
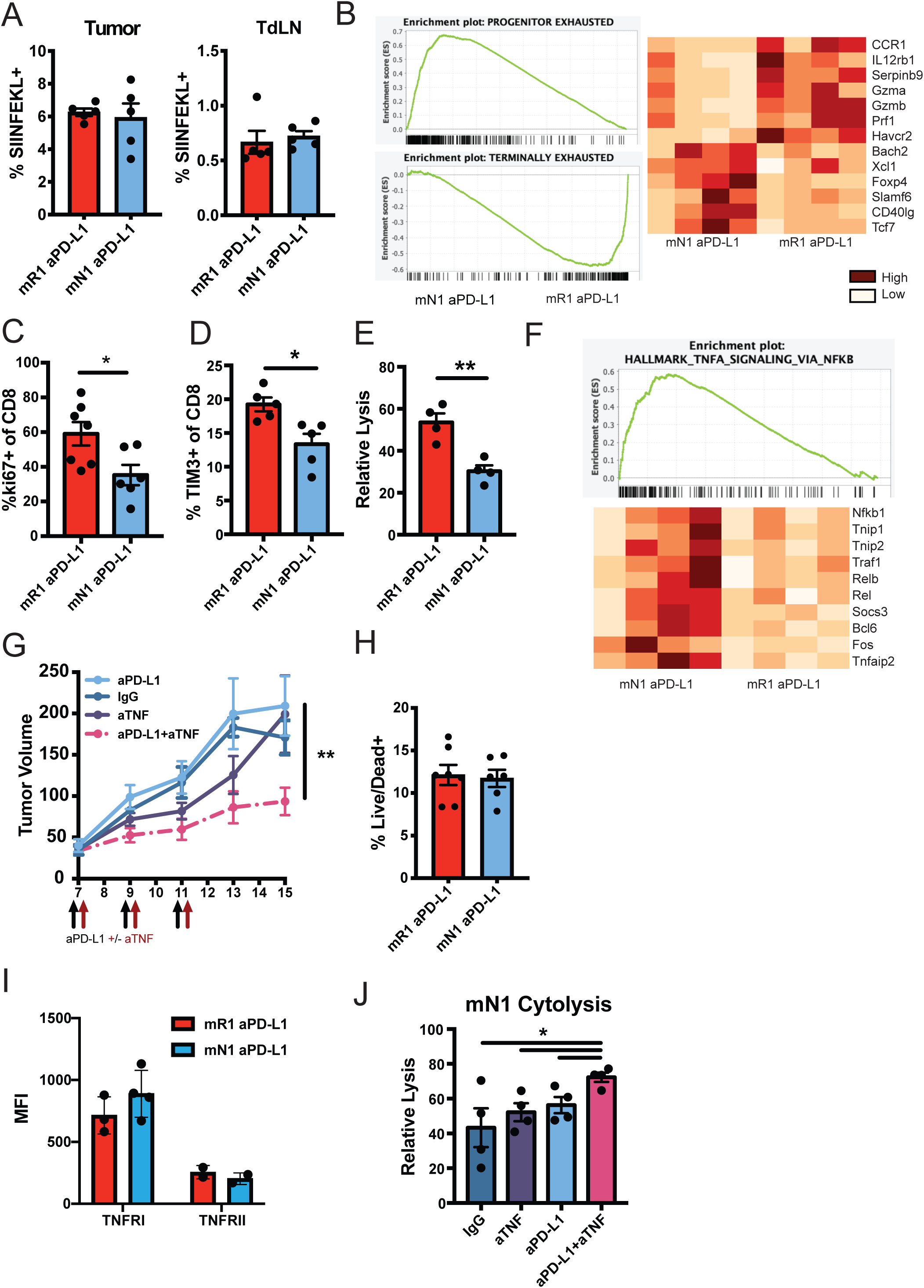
Non-Responder Microbiota Reduce Tumor Effector CD8^+^ Response. (A) Frequency of SIINFEKL-positive cells in the tumor and draining lymph node at day 10 of B16-ova tumor growth, mean ± SEM. (B) CD8^+^ T cells were sorted from the tumors of anti-PD-L1-treated mR1 or mN1 mice at day 10 and the RNA was sequenced. Libraries were compared to gene sets for stem-like or effector-like exhausted T cells. Representative heatmap of the top genes in replicate mice for each signature is shown. Relative expression compared to mean for all samples, max=3.4, min=0.26 (C) Frequency of tumor CD8 T cells expressing Ki-67, mean ± SEM. (D) Tumor CD8 T cells expressing TIM-3, mean ± SEM. (E) SIINFEKL loaded target cells were injected into B16-ova bearing mice. Live cell frequency was analyzed in the lymph node 24 hours later. Data representative of three independent experiments. (F) Gene set enrichment for Hallmark TNF Receptor signaling. Relative expression heatmap compared to mean for all samples in a row; max=5.03, min=0.49. (G) Tumor growth with TNF blockade in mice colonized with the mN1 nonresponder community, mean ± SEM (H) Frequency of dead cells in tumor cell suspensions from day 10, mean ± SEM. (I) MFI of TNFRI and TNFRII, mean ± SEM. (J) Frequency of killed SIINFEKL+ cells in B16-ova tumors treated with anti-TNF, mean ± SEM. All data analyzed at day 10 of tumor growth. *p<0.05, **p<0.01.

Similar to chronic infection, tumors are characterized by the generation of exhausted T cells, which transcriptionally differ from traditional memory and effector T cells [30]. Single-cell sequencing studies have demonstrated that exhausted T cells can be subtyped as either stem-like exhausted cells or terminal effector cells [31]. While stem-like cells have little cytotoxic capacity, they produce more cytokines and can differentiate into effector cells that adopt cytotoxicity [6]. The presence of the stem-like cells is necessary for durable response to checkpoint blockade, and these cells proliferate and differentiate into terminal effector cells after PD-1 blockade [7]. To profile the CD8^+^ T cell phenotype, we performed RNA sequencing of CD8^+^ TILs in anti-PD-L1 treated mN1 and mR1 mice. Gene Set Enrichment Analysis [32] showed stem-like associated genes [6] in the mN1 colonized animals, which contrasted with enrichment of effector-associated genes in mR1 tumors (fig. 3B). Supporting the expansion of effector cells in mR1 responders, TILs had higher proliferation measured by Ki-67 staining (fig. 3C) and a higher frequency of the effector marker Tim-3 (fig. 3D). Expanding on results in the tumor, we found increased expression of the stem-like marker Slamf6 in T cells of the tumor-draining lymph node in mN1 mice (fig. S3F), and reduced expression of Ki-67 by these cells (fig. S3G). This data supports a model of reduced tumoral expansion of effector T cells.

Immune mediated control of tumor growth is ultimately determined by the ability of T cells to kill tumor cells. To evaluate antigen specific cytotoxicity, mice bearing B16-ova tumors were subjected to *in vivo* cytolysis assays. When we measured survival of i.v. injected, SIINFEKL-loaded target splenocytes, mN1 mice killed significantly fewer SIINFEKL^+^ target cells than mR1 mice, further supporting expansion of functional effector T cells in mR1 responders (fig. 3E). Finally, we examined cytokine production by CD8^+^ T cells and found a significantly higher frequency of cells in the tumor able to produce both interferon-γ and TNFα after PMA/Ionomycin stimulation (figs. S3H) and draining lymph node (fig. S3J). This result further demonstrates enrichment for stem-like exhausted cells in mN1 mice because stem-like cells have been shown to produce both TNFα and IFNγ *in vitro* and *in vivo* while effector-like cells are restricted to IFNγ production [6]. These results show that the microbiota can influence the balance of exhausted T cell subsets, important for effective immunotherapy response.

Elevated TNFα has previously been shown to limit the efficacy of PD-1 blockade [33, 34], so we were interested in whether the increased TNFα found in mN1 mice had an effect on T cell phenotype and function. Suggesting its importance, CD8^+^ TILs from mN1 mice showed enrichment of genes downstream of TNFα signaling, notably NF-kB, RELB, TRAF1, TNAIP3, and TRAF2 (fig. 3F). To evaluate the clinical role of TNFα signaling, we treated mN1 mice with anti-PD-L1 in combination with anti-TNFα. While blockade of TNFα alone had no effect on tumor growth, combination with anti-PD-L1 reduced tumor growth in these gnotobiotic mice (fig. 3G). The negative effects of TNFα on PD-1 therapy have been previously attributed to increased apoptosis of CD8^+^ T cells in the tumor microenvironment (Bertrand et al., 2017). In contrast to those reports, we did not find any difference in CD8^+^ T cell death in the tumors of anti-PD-L1 treated mN1 and mR1 mice by live/dead staining (fig. 3H). Apoptosis of CD8^+^ T cells in this previous work relies on death domain-containing TNFRI. TNFRII by contrast does not contain a death domain and is thought to work instead through suppression of T cell activity [35]. Signaling through TNF receptors has been shown to reduce their respective surface expression as the result of receptor internalization [36]. While TNFRI had much higher expression than TNFRII on tumor CD8^+^ T cells, we did not find significant differences in these receptors between mN1 and mR1 mice (fig. 3I). However we did find that mice on dual PD-L1/TNF blockade showed increased antigen specific cytolysis, which supports a role for TNFα in constraining the effector T cell response in anti-PD-L1-treated mN1 mice (fig. 3J). Taken together these results highlight that microbiota-induced anti-PD-L1 failure can be overcome by TNFα blockade that increases cytotoxicity against the tumor. They also highlight a novel potential strategy to overcome checkpoint blockade failure in patients.

### Microbial Transplant Enhances the Effector CD8^+^ T Cell Phenotype and Reverses anti-PD-L1 Failure in mN1 Mice

Given the potential of gut microbiota alterations to modify immune activity [14, 18, 37], we investigated if a defined microbiota transplant would render tumors in mN1 mice susceptible to PD-L1 blockade. Gnotobiotic mice were colonized with mN1 microbiota for 4 weeks before oral gavage of the mR1 community, either 2 days before tumor injection or 2 days before anti-PD-L1 therapy (fig. S4A). In both cases, mR1 transplanted mice showed reduction of tumor growth with anti-PD-L1 (fig. 4A). Metagenomic sequencing verified successful engraftment of the mR1 microbiota, comprising ∼20-30% of relative abundance (figs 4B, S4B, and S4C). When we examined the specific community members that were modified, we found significant enrichment of donor *Escherichia coli* (4.6%) and *Ruminococcus gnavus* (7.3%), as well as members of the *Bacteroides* genus, *B. fragilis* (5.5%) and *B. ovatus* (3.5%) (fig. S4D). Of note, *Bifidobacterium* species constituted 7% of the fecal microbiome of donor mice, but they represented 0.44% in the FMT mice (fig. S4E). This finding was interesting given the role previously demonstrated for *Bifidobacterium* in microbiota-induced anti-PD-1/PD-L1 response [12, 23]. The expansion of these taxa post-FMT was paralleled by reduction of *B. uniformis* and *Ruminococcus albus* (fig. S4F). The composition of *R. albus* was reduced to barely detectable levels in all transplanted mice.

**Figure 4.**
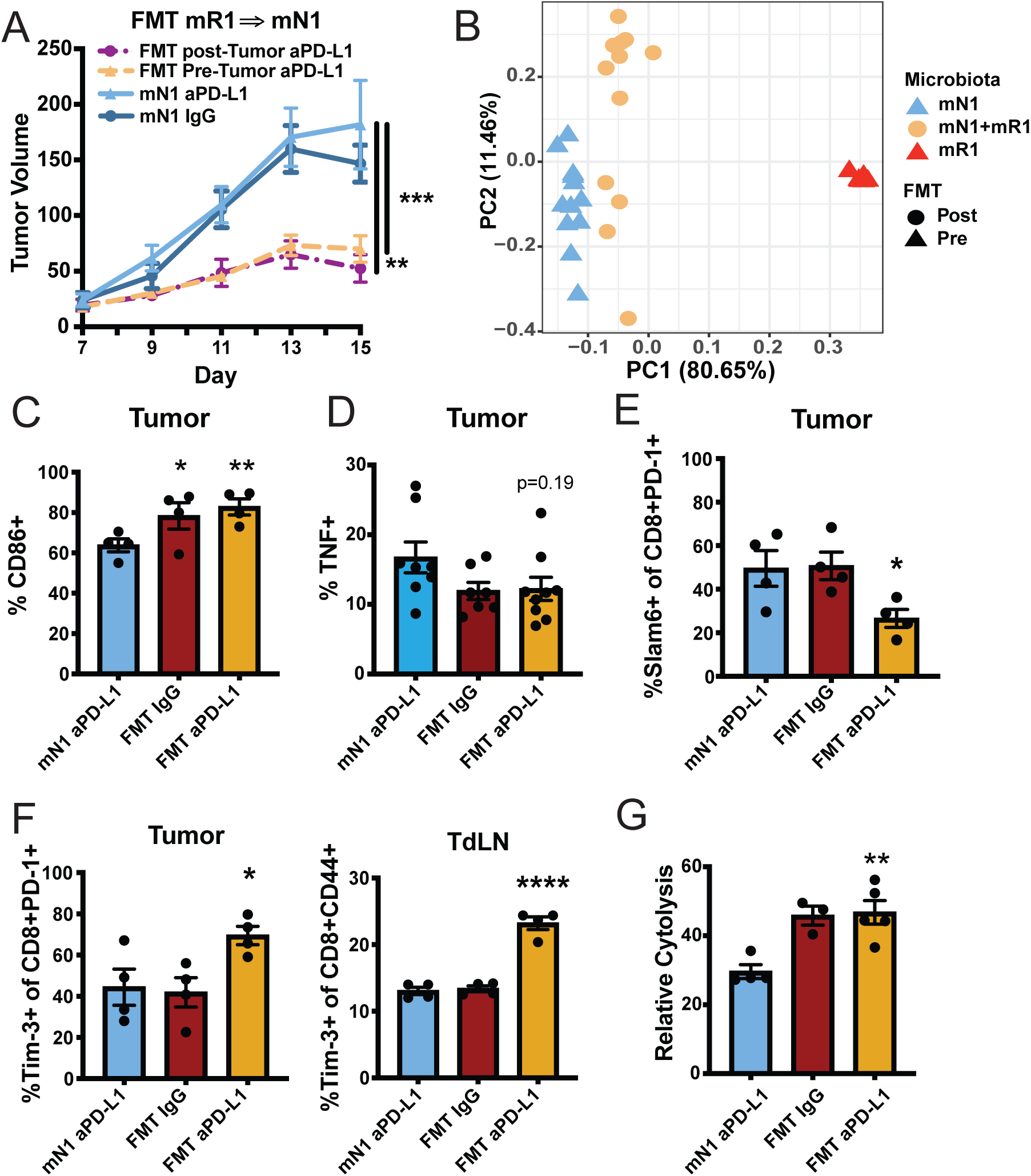
Microbial Transplant Restores Anti-PD-L1 Response and Cytotoxicity in Non-Responder Gnotobiotic Mice. (A) mN1 mice were colonized for 3 weeks before oral gavage of mR1 bacterial community either 2 days before tumor injection (Pre-Tumor) or 2 days before anti-PD-L1 was initiated (Post-Tumor). Data summary of two independent experiments. (B) Fecal samples were collected at sacrifice and assessed for community composition by metagenomic sequencing. Composition compared to mR1 fecal samples from a separate experiment. (C) CD86 expression on cDC1, mean ± SEM. (D) TNF production by tumor cDC1, mean ± SEM. (E) Frequency of slamf6^+^ cells in the exhausted PD-1^+^ T cell pool, mean ± SEM. (F) Frequency of TIM-3 positive cells in the tumor and tumor draining lymph node, mean ± SEM. (G) Frequency of killed SIINFEKL+ cells in B16-ova tumors. Data summary of two independent experiments. *p<0.05, **p<0.01, ***p<0.001, ****p<0.0001

To understand the role of the microbiota in driving immune modification we assessed changes within the tumor and draining lymph node. Transplanted animals, similar to mR1 singly colonized animals, had higher expression of CD86 on dendritic cells (fig. 4C), and a trend toward reduced TNFα production (fig. 4D). Evaluating the adaptive immune response, we also saw that mR1 transplantation reduced the proportion of stem-like cells measured by Slamf6 positivity (fig. 4E), and increased the frequency of effector-like T cells, measured by Tim-3 expression, in both the tumor and draining lymph node (fig. 4F) The phenotype was functionally verified by increased killing of SIINFEKL^+^ target cells after FMT (fig. 4G). Interestingly we also found that the frequency of cDC1 presenting SIINFEKL on MHC class I was significantly elevated in transplanted mice (fig. S4G). These results are consistent with a role for FMT in regulating exhausted T cell differentiation in the context of PD-L1 blockade, while improving clinical outcome.

## Discussion

We have shown that the gut microbiota is capable of inhibiting response of B16 melanoma to anti-PD-L1, likely through effects on immune recruitment and activity of both APCs and CD8^+^ T cells. Specifically, the microbiota regulates the balance of exhausted CD8^+^ T cell subtypes in the tumor microenvironment, and this effect can be overcome by either microbial transplant or concomitant blockade of TNFα. Several studies have demonstrated that the exhaustion transcriptional signature of T cells subjected to chronic antigen exposure in either the chronic infection setting or tumor microenvironment is driven by the transcription factor Tox and mediated by PD-1 [38]. Within this pool of exhausted cells, chromatin accessibility regions have been defined for progenitor-like cells, characterized by Slamf6 and Tcf-1 driven transcription, while terminal effectors are characterized by high levels of granzymes and coinhibitory receptors like Tim-3 [39]. PD-1 blockade drives proliferation of this exhausted T cell pool [40]. Still unclear are the signals that regulate the balance of progenitor and effector exhausted cells. Regulation by the microbiota is a particularly interesting target, given the myriad influences of commensal microbes on local and systemic immune regulation [41].

We were surprised to find that TNFα production did not mirror costimulatory expression in antigen presenting cells. Typically, both proinflammatory cytokines and costimulatory molecules are increased during APC maturation. There is however an interesting role for TNFα in suppression of immune responses and immune exhaustion. One of the first discoveries on HIV was that viral load positively correlated with serum TNFα level, suggesting TNFα might drive impaired viral control [42]. Much work has been done to characterize the dysfunctional biology of exhausted T cells by studying chronic infection using mouse lymphocytic choriomeningitis virus (LCMV). In particular, focus has been on LCVM-c13, which responds to PD-1/PD-L1 blockade with viral clearance [43]. LCMV-WE, like LCMV-c13 infection, is characterized by the presence of phenotypically exhausted T cells, however mice with chronic LCMV-WE also demonstrate persistently elevated serum TNFα [44]. When infected mice were treated with anti-TNFα, they were able to clear the virus [45]. This result highlights the possibility of the TNF-axis in the regulation of T cell exhaustion. Finally, PD-1 blockade works partly by preventing the PD-1/Shp-2 mediated inactivation of CD28 [3, 4]. Interestingly, TNFα exposure directly downregulates CD28 in vitro, and rheumatoid arthritis patients, in the presence of high serum TNFα concentrations, develop CD28 negative T cells with reduced function [46] – what today might be called exhausted T cells.

The results presented here highlight the importance of the microbiota in negatively regulating anti-tumor responses to PD-L1 blockade. While there are several associations between TNFα and T cell function in chronic antigen exposure, the mechanism through which TNFα may regulate exhaustion remains hidden. Moreover, the route through which the microbiota modifies immune activity at the tumor site is yet unknown. In the most evocative work to date, Tanoue et al. suggested that microbial metabolites were able to direct immune tone and anti-PD-1 response in tumor bearing mice, but a causal role for metabolites remains elusive [14]. The discovery of different immune activation status in tissues of tumor-naïve mN1 and mR1 mice highlights the complex role of microbial signals in directing the immune response. Taken together, this work highlights the role of the microbiota as an inhibitor of checkpoint blockade immunotherapy responses and suggests avenues for better understanding T cell function in the tumor microenvironment.

## Methods

### Mice

All animal experiments performed in this study were approved by the Institutional Animal Care and Use Committee (IACUC) of Icahn School of Medicine at Mount Sinai. Specific pathogen free (SPF) mice were purchased from Jackson Labs or Charles River. Germfree (GF) WT C57BL/6 mice were housed in standard, commercially available flexible vinyl isolators. To facilitate high-throughput gnotobiotic studies, we utilized previously described “out-of-the-isolator” gnotobiotic techniques [17, 47]. Gnotobiotic mice were generated by colonizing GF mice with defined cultured human microbial communities at 4-6 weeks old [37]. Under strict aseptic conditions, germfree mice were transferred to autoclaved filter-top cages outside the breeding isolator and colonized with human microbiota by a single oral gavage of 200ul of previously frozen bacterial cocktail in glycerol. All experiments were conducted at least 21 days after colonization. As standard in the community, all tumor experiments were performed in female mice, owing to behavior amenable for flank tumor growth.

### Cultured Microbial Communities

To generate arrayed culture collections from human donors, clarified and diluted donor stool was plated onto a variety of solid selective and non-selective media under anaerobic, micro-aerophilic and aerobic conditions. Plates were incubated for 48-72 hours at 37C. 384 single colonies from each donor microbiota were individually picked and grown in liquid LYBHIv4 media for 48 hours under anaerobic conditions. Regrown isolates were identified at the species level using a combination of MALDI-TOF mass spectrometry (Bruker Bioytyper) and 16S rRNA amplicon sequencing. Unique Isolates are reported in supplementary table 1. All regrown isolates were pooled by equal volume and stored in LYBHIv4 media with 15% glycerol at −80C for administration to gnotobiotic mice.

### Tumor Models

The C57BL/6-derived melanoma cell line B16-F1, the B16-OVA-transfected clone and the B16-YFP-transfected clone were maintained at 37C with 5% CO_2_ in RPMI+10%FBS+1%pen/strep. 3 × 10^5^ B16-F1 cells, or 5 × 10^5^ B16-OVA/B16-YFP cells were injected subcutaneously in the flank in 50uL PBS. Tumor size was determined by the formula: Length × Width^2^ × 0.5 on the indicated days. Tumor volume ratios were calculated as: log_2_ (*TreatedTumorVolume*_*i*_ /*UntreatedTumorVolume*_*i*_), where *TreatedTumorVolume* is the median tumor size on day *i* for mice treated with anti-PD-L1 and *UntreatedTumorVolume* is the median tumor size on day *i* for mice treated with IgG isotype control.

### Treatments

Mice were initiated on anti-PD-L1 treatment (200ug/mouse per dose; 10F.9G2, BioXcell) or isotype (rat IgG2b, BioXcell) by i.p. injection on days 7, 9, and 11 of tumor growth. In some experiments, anti-TNFα (200ug/dose; XT3.11, BioXcell) or IgG1 isotype (BioXcell) was administered on days 7, 9, 11, and 13. Antibodies were injected in 100ul PBS. In some experiments mice were treated with an antibiotic cocktail of metronidazole (1mg/ml), ampicillin (1mg/ml), neomycin (1mg/ml), and vancomycin (0.5mg/ml) by drinking water. In microbial transplant experiments, mice were gavaged on the given day with 200ul of the donor bacteria library in glycerol.

### Cell Preparation

Single cell suspensions of the tumor were obtained after tumor digestion with 400U/mL of collagenase D (Roche) at 37C for 1 hour. In some experiments, hematopoietic cells were enriched by density gradient centrifugation with 40/90 Percoll gradient (GE Healthcare Life Sciences) for 30 min at 2,500 rpm. To obtain DCs, lymph nodes were digested in 400U/mL collagenase D for 45 min at 37C. When only lymphocytes were required, the lymph node was minced and passed through a 70um filter. To isolate cells from the lamina propria, the colon was harvested, cleared of food debris, transected longitudinally, and a razor blade was used to clear the mucus layer from the tissue. The colon sample was then incubated in dissociation buffer (HBSS without calcium and magnesium, 5% FBS, 5mM EDTA, and 15mM HEPES) for 30 minutes, before incubation in 0.5mg/mL Collagenase IV and 0.5mg/mL DNase for 1 hour at 37C. To isolate cells from the skin, the ear was excised from the mouse and separated into dorsal and ventral sheets. The sheets were placed dermis-side down in HBSS containing 4U/mL dispase for 90 min at 37C. The dermis and epidermis were then separated, washed in PBS, and minced with scissors. The tissue was digested in collagenase D and DNase for 45 min at 37C. In all experiments, tissue chunks were ground through a 70um filter using the plunger of a 3mL syringe.

### Flow Cytometry

Cells were labeled with the antibodies in Supplementary Table 4. In some experiments, cells were stimulated with 100ng/mL PMA (Sigma) and 0.5ug/mL ionomycin (Sigma-Aldrich) at 37C for 3 hr in 10ug/mL Brefeldin A (Sigma) to allow accumulation of cytokines. When measuring cytokines in myeloid cells, cells were rested in Brefeldin A for 3 hours at 37C without any ex vivo stimulation. After staining for surface markers, cells were fixed and permeabilized, followed by staining for IFN-γ, TNFα, IL-1b, or IL-12. Antigen-specific T cell generation was determined by staining with MHCI(H-2Kb)-SIINFEKL tetramers obtained from the NIH Tetramer Core Facility (https://tetramer.yerkes.emory.edu/).

### In Vivo Cytolysis

To measure antigen-specific killing in gnotobiotic mice, 500,000 B16-ova tumor cells were injected subcutaneously in the flank. On day 7, mice were treated with anti-PD-L1 or control. On day 9, WT SPF mice were sacrificed and splenocytes harvested by passing tissue through a 70um filter. The splenocytes were RBC lysed before incubation with 1uM or 10nM SIINFEKL peptide in RPMI+10%FBS+1% penicillin/streptomycin and 25uM B-mercapthoethanol for 1 hour at 37C. Peptide loaded cells were then incubated in 50uM or 5uM CFSE in PBS for 10 minutes at RT. Differentially loaded and labeled cells were mixed, then 5*10^5^ cells were intravenously administered in 300uL through retro-orbital injection. Mice were given second dose of anti-PD-L1 and/or anti-TNFα at this time. 20-26 hours later, mice were sacrificed. The tumor draining lymph node was isolated and digested in collagenase for 30 minutes at 37C. A shorter digestion time is sufficient to collect T cells from the lymph node. Cells were then acquired on a BD Fortessa, and the ratio of CFSEhi/CFSElo in gnotobiotic mice was compared to that of tumor-naïve SPF mice to calculate % specific lysis as previously described [21, 48].

### RNA Sequencing

Tumors were digested, stained for surface antibodies and sorted on the BD-FACSAria II (BD Biosciences) into Trizol for RNA extraction. Sorted cells were CD45^+^CD3^+^CD8^+^Live-dead^neg^. cDNA libraries were prepared with SmartSeq v4 Ultra Low Input RNA Kit for Sequencing (Takara). cDNA was barcoded and pooled equimolar for sequencing on Illumina MiSeq (Single-end 150bp). Transcript abundances were quantified using the Ensembl GRCm38 cDNA reference using Kallisto version 0.44.0. Transcript abundances were summarized to gene level using tximport. Expression matrices were filtered to above 10 tpm. Gene Set Enrichment Analysis was performed using GSEA 4.0.1 (Broad Institute) using ‘gene_set’ permutation type. Progenitor-like and Terminally Differentiated genesets were constructed from the significant differentially regulated genes in [6].

### Metagenomics

Mouse fecal samples were subjected to a protocol for DNA isolation through bead beating [49]. Briefly, 200uL of 0.1mm diameter zirconia/silica beads and 700uL of DNase inactivation buffer (2.5g SDS, 500 uL 0.5M EDTA, 5mL 1M Tris, 494.5mL H_2_O) were added to the fecal samples. Tubes were capped and bead-beat for 5 minutes. DNA was then isolated using QuiaQuick (Quiagen) columns and quantified by the Qubit assay (Life Technologies). Sequencing libraries were generated from sonicated DNA with the NEBNext Ultra II DNA Library Prep Kit (New England BioLabs). Ligation products were purified with SPRIselect beads (Beckman Coulter) and enrichment PCR performed with NEBNext Ultra Q5 Master Mix (New England BioLabs). Samples were pooled in equal proportions and size-selected using 0.6x then 0.2x AMPure beads (Beckman Coulter) before sequencing with an Illumina HiSeq (paired-end 150bp). Reads from metagenomic samples were trimmed, subsampled to 100,000 reads and mapped to the unique regions of bacterial genomes known to potentially form part of the microbiota in a given gnotobiotic sample. Abundances were scaled to the size of each specific genome as previously described [50]. Principle component analysis was calculated using the ggfortify R package (https://cran.r-project.org/web/packages/ggfortify/ggfortify.pdf) and ‘prcomp’ function on relative abundance at the species level.

## Tables

**Supplementary Table 1** | **Cultured Human Microbiota Composition**.

**Supplementary Table 2** | **Pairwise Microbial Comparisons**

**Supplementary Table 3** | **Shared Bacterial Species**

**Supplementary Table 4** | **Antibodies**

---

**Supplement 1.**
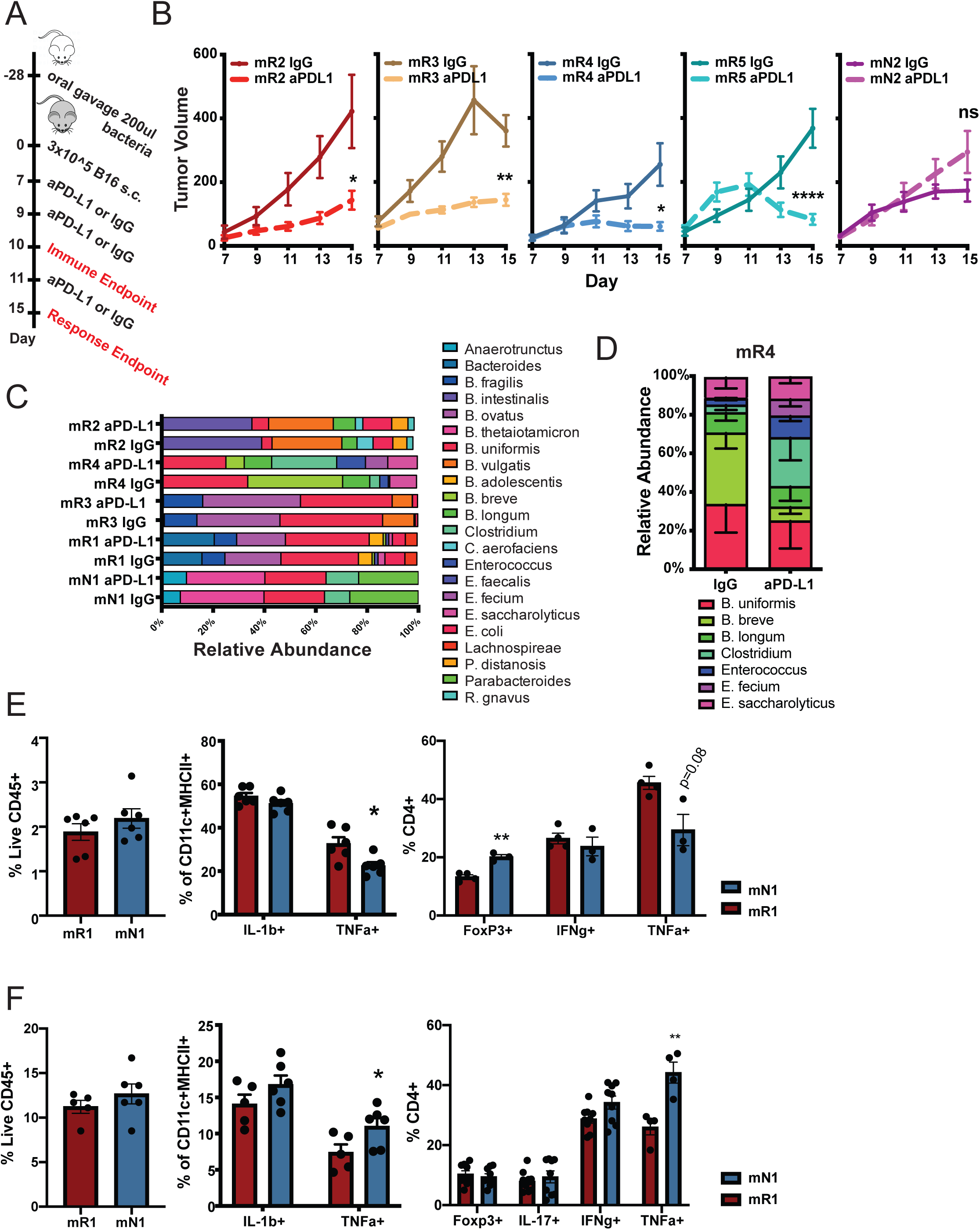
Microbiome Regulates Immune Tone but Does Not Change with Anti-PD-L1. (A) Overview of anti-PD-L1 treatment and analysis of gnotobiotic tumor growth. (B) Tumor growth plot for additional microbiota, data represent mean ± SEM for at least two experiments. (C) Microbiota composition at the species level was compared for the five communities for which we have complete genomes. Two-way ANOVA insignificant for all but the mR4 microbiota. (D) mR4 microbial abundance at the species level, mean ± SEM. (E) Cytokine production by colon lamina propria APCs or CD4 T cells in the mesenteric lymph node, mean ± SEM. (F) Cytokine production by skin APCs or CD4 cells in the skin draining lymph node, mean ± SEM. Data representative of two independent experiments. *p<0.05, **p<0.01, ***p<0.001, ****p<0.0001

**Supplement 2.**
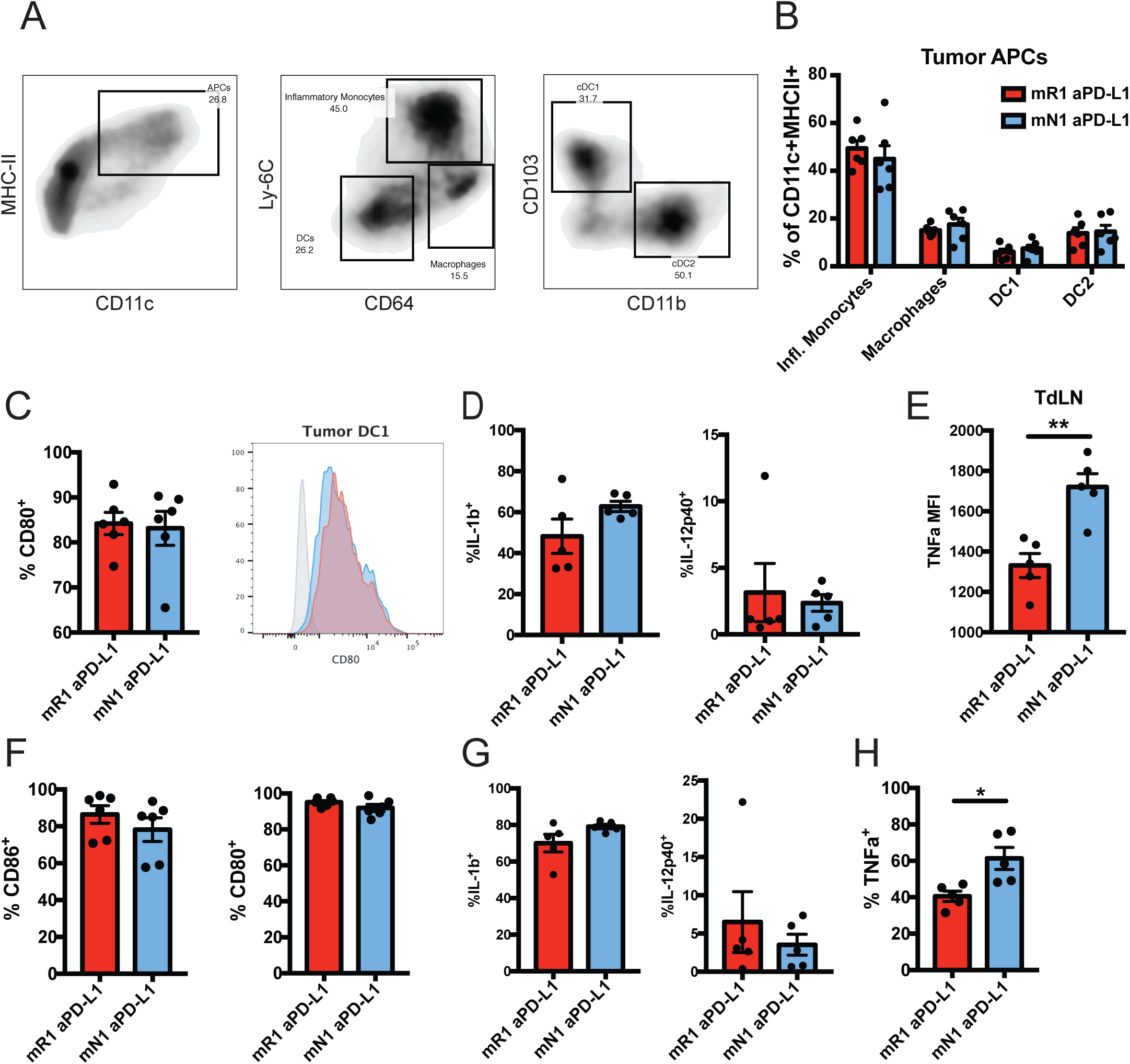
Increased TNFα Production by Tumor DCs and Macrophages in mN1 mice. (A) Gating used to identify antigen presenting cells of the tumor. (B) Breakdown of CD11c^+^MHCII^+^ antigen presenting cells at day 10, mean ± SEM. (C) CD80 expression on tumor cDC1, mean ± SEM. (D) Resting production of IL-1b or IL-12 in cDC1 from tumors at day 10, mean ± SEM. (E) Expression of TNF by migratory cDC1 in the draining lymph node. All migratory cDC1 were TNF^+^. (F) Resting CD86 and CD80 expression of tumor macrophages (CD64^+^Ly6C^−^), mean ± SEM. (G) Resting production IL-1b and IL-12 tumor macrophages (H) Macrophage production of TNFα, mean ± SEM. All data collected at day 10 during anti-PD-L1 treatment. *p<0.05, **p<0.01.

**Supplement 3.**
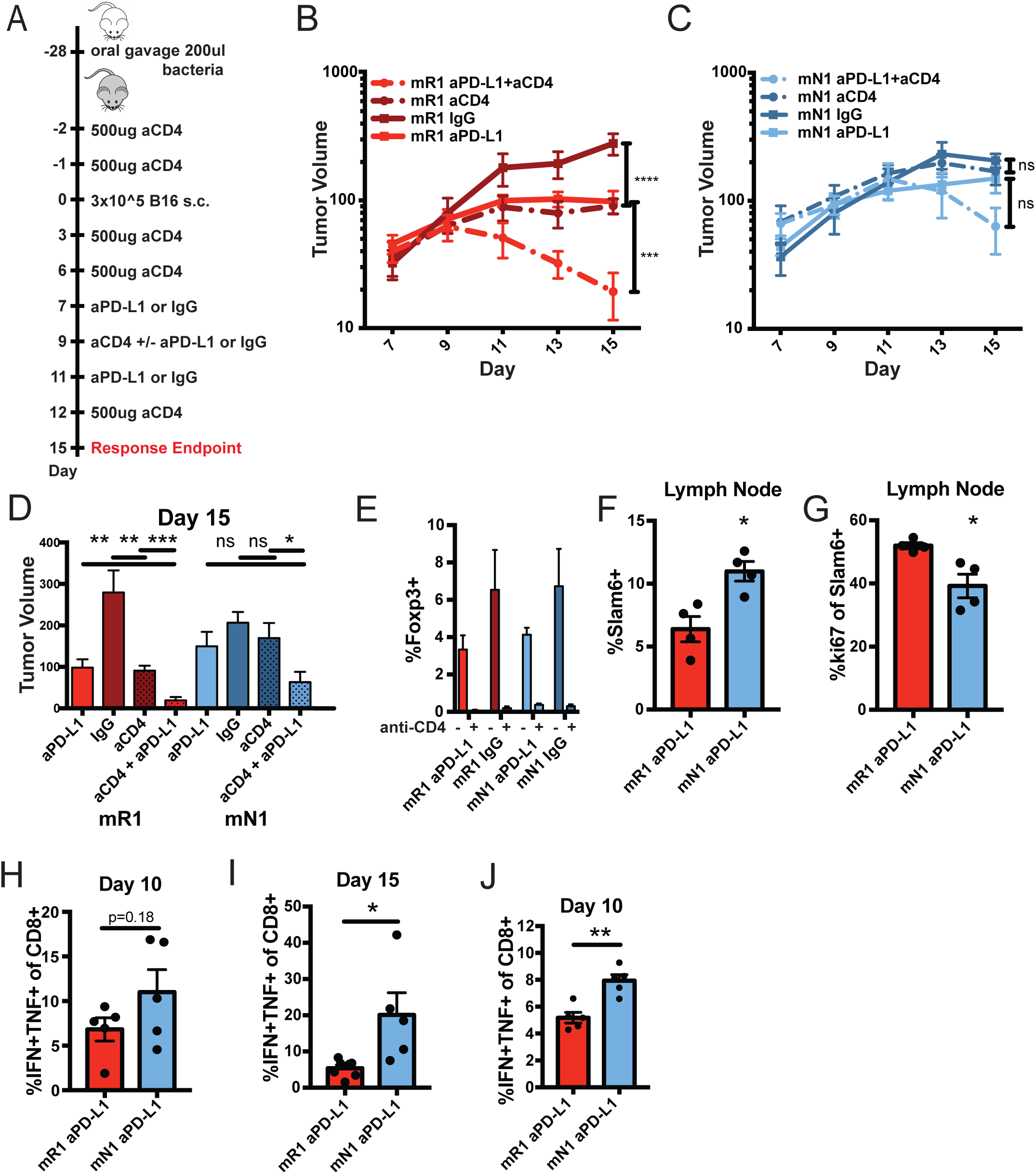
CD4 Depletion Does Not Rescue anti-PD-L1 Response in Non-Responder Gnotobiotic Mice. (A) Schematic for experimental depletion of CD4 cells. (B) Tumor growth with or without CD4 depletion in mR1 responders. (C) Tumor growth with or without CD4 depletion in mN1 non-responders. Data summary of two independent experiments in head-to-head comparison with (B). (D) Mean + SEM for tumor volume at sacrifice. (E) T_reg_ frequency with or without CD4 depletion at sacrifice. (F) Frequency of Slamf6 positive (stem-like) cells in the tumor draining lymph node. (G) Ki-67 expression of slamf6+ cells in the tumor draining lymph node. (H) Frequency of IFNγ^+^TNFα ^+^ CD8 T cells in the tumor at day 10 and (I) day 15, mean ± SEM. (J) Frequency of IFNγ^+^TNFα ^+^ CD8 T cells in the tumor draining lymph node at day 10, mean ± SEM. *p<0.05, **p<0.01, ***p<0.001

**Supplement 4.**
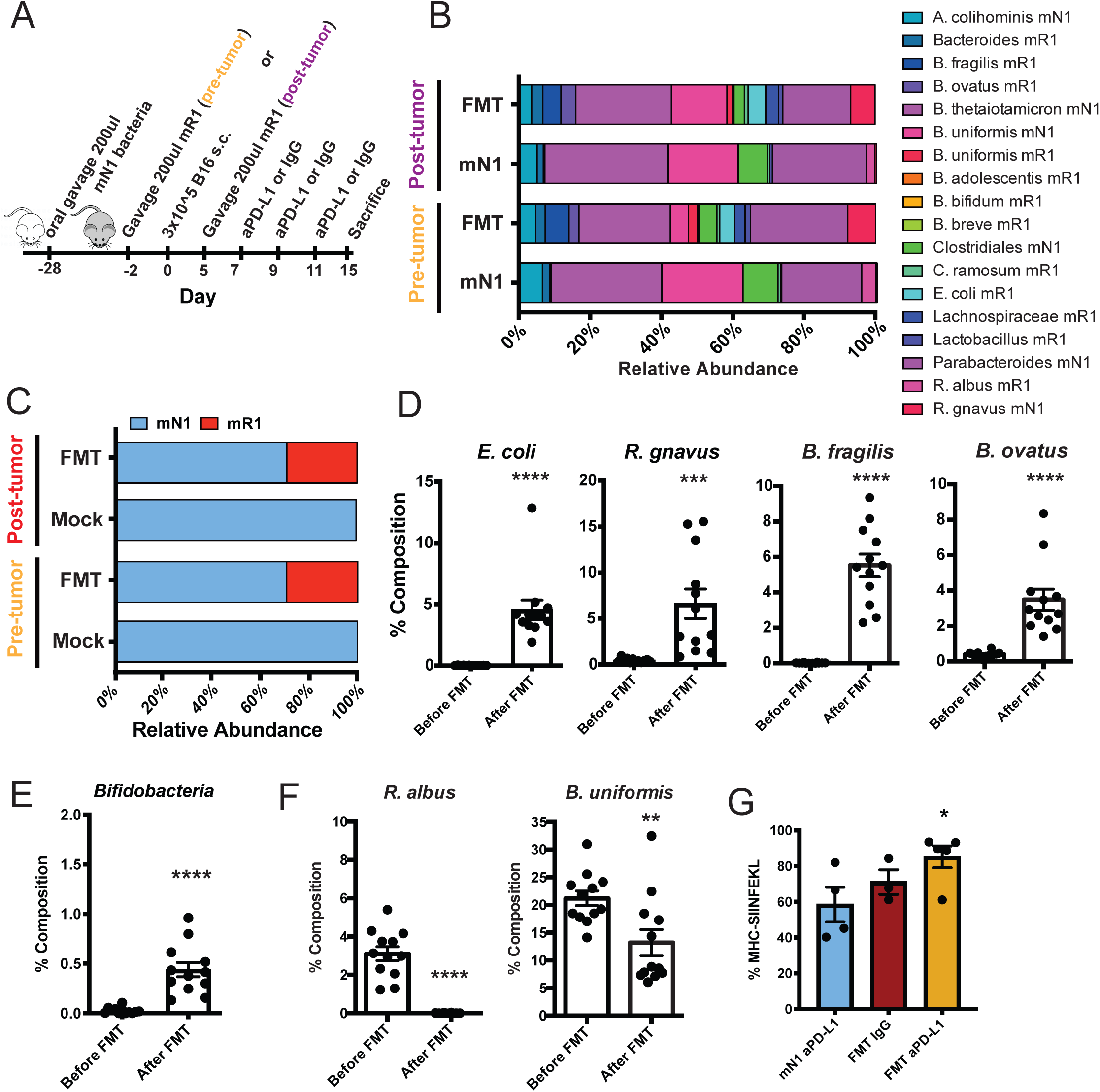
Engraftment of Responder Microbes into Non-Responder Mice. (A) Schematic of microbial transplant (B) Fecal microbiota abundance with and without FMT at the species level. (C) Percentage species abundance in fecal microbiota by donor and recipient in transplanted animals. (D) Abundance of highly engrafted species after transplantation, mean ± SEM. (E) Relative abundance of strains in the *Bifidobacterium* genus, mean ± SEM. (F) Highly reduced microbes in the recipient after transplantation, mean ± SEM. (G) Expression of SIINFEKL-MHC class I on tumor cDC1 cells. *p<0.05, **p<0.01, ***p<0.001, ****p<0.0001

## Notes

### Competing Interest Statement

The authors have declared no competing interest.

## References

1. Baumeister, S.H., et al., Coinhibitory Pathways in Immunotherapy for Cancer. Annual Review of Immunology, Vol 34, 2016. 34: p. 539–573.

2. Postow, M.A., et al., Nivolumab and ipilimumab versus ipilimumab in untreated melanoma. N Engl J Med, 2015. 372(21): p. 2006–17.

3. Kamphorst, A.O., et al., Rescue of exhausted CD8 T cells by PD-1-targeted therapies is CD28-dependent. Science, 2017. 355(6332): p. 1423–1427.

4. Hui, E.F., et al., T cell costimulatory receptor CD28 is a primary target for PD-1-mediated inhibition. Science, 2017. 355(6332): p. 1428-+.

5. Wherry, E.J. and M. Kurachi, Molecular and cellular insights into T cell exhaustion. Nat Rev Immunol, 2015. 15(8): p. 486–99.

6. Miller, B.C., et al., Subsets of exhausted CD8(+) T cells differentially mediate tumor control and respond to checkpoint blockade. Nat Immunol, 2019. 20(3): p. 326–336.

7. Jadhav, R.R., et al., Epigenetic signature of PD-1+TCF1+CD8 T cells that act as resource cells during chronic viral infection and respond to PD-1 blockade. Proceedings of the National Academy of Sciences of the United States of America, 2019. 116(28): p. 14113–14118.

8. Zou, W., J.D. Wolchok, and L. Chen, PD-L1 (B7-H1) and PD-1 pathway blockade for cancer therapy: Mechanisms, response biomarkers, and combinations. Sci Transl Med, 2016. 8(328): p. 328rv4.

9. Swaika, A., W.A. Hammond, and R.W. Joseph, Current state of anti-PD-L1 and anti-PD-1 agents in cancer therapy. Mol Immunol, 2015. 67(2 Pt A): p. 4–17.

10. Michot, J.M., et al., Immune-related adverse events with immune checkpoint blockade: a comprehensive review. Eur J Cancer, 2016. 54: p. 139–148.

11. Belkaid, Y. and T.W. Hand, Role of the microbiota in immunity and inflammation. Cell, 2014. 157(1): p. 121–41.

12. Matson, V., et al., The commensal microbiome is associated with anti-PD-1 efficacy in metastatic melanoma patients. Science, 2018. 359(6371): p. 104-+.

13. Routy, B., et al., Gut microbiome influences efficacy of PD-1-based immunotherapy against epithelial tumors. Science, 2018. 359(6371): p. 91–97.

14. Tanoue, T., et al., A defined commensal consortium elicits CD8 T cells and anti-cancer immunity. Nature, 2019. 565.

15. Gopalakrishnan, V., et al., Gut microbiome modulates response to anti-PD-1 immunotherapy in melanoma patients. Science, 2018. 359(6371): p. 97–103.

16. Gopalakrishnan, V., et al., The Influence of the Gut Microbiome on Cancer, Immunity, and Cancer Immunotherapy. Cancer Cell, 2018. 33(4): p. 570–580.

17. Britton, G.J., et al., Microbiotas from Humans with Inflammatory Bowel Disease Alter the Balance of Gut Th17 and RORgammat(+) Regulatory T Cells and Exacerbate Colitis in Mice. Immunity, 2019. 50(1): p. 212–224 e4.

18. Yang, C., et al., Fecal IgA Levels Are Determined by Strain-Level Differences in Bacteroides ovatus and Are Modifiable by Gut Microbiota Manipulation. Cell Host Microbe, 2020. 27(3): p. 467–475 e6.

19. Vetizou, M., et al., Anticancer immunotherapy by CTLA-4 blockade relies on the gut microbiota. Science, 2015. 350(6264): p. 1079–84.

20. Moynihan, K.D.e.a., Eradication of large established tumors in mice by combination immunotherapy that engages innate and adaptive immune responses. Nature Medicine, 2016. 5(3): p. 269–275.

21. Salmon, H., et al., Expansion and Activation of CD103(+) Dendritic Cell Progenitors at the Tumor Site Enhances Tumor Responses to Therapeutic PD-L1 and BRAF Inhibition. Immunity, 2016. 44(4): p. 924–38.

22. Tumeh, P.C., et al., PD-1 blockade induces responses by inhibiting adaptive immune resistance. Nature, 2014. 515(7528): p. 568–71.

23. Sivan, A., et al., Commensal Bifidobacterium promotes antitumor immunity and facilitates anti-PD-L1 efficacy. Science, 2015. 350(6264): p. 1084–9.

24. Hammerich, L., et al., Systemic clinical tumor regressions and potentiation of PD1 blockade with in situ vaccination. Nat Med, 2019. 25(5): p. 814–824.

25. Hildner, K., et al., Batf3 deficiency reveals a critical role for CD8alpha+ dendritic cells in cytotoxic T cell immunity. Science, 2008. 322(5904): p. 1097–100.

26. Maier, B., et al., A conserved dendritic-cell regulatory program limits antitumour immunity. Nature, 2020. 580(7802): p. 257–262.

27. Garris, C.S., et al., Successful Anti-PD-1 Cancer Immunotherapy Requires T Cell-Dendritic Cell Crosstalk Involving the Cytokines IFN-gamma and IL-12. Immunity, 2018. 49(6): p. 1148–1161 e7.

28. Huang, A.C., et al., T-cell invigoration to tumour burden ratio associated with anti-PD-1 response. Nature, 2017. 545(7652): p. 60–65.

29. Ueha, S., et al., Robust Antitumor Effects of Combined Anti-CD4-Depleting Antibody and Anti-PD-1/PD-L1 Immune Checkpoint Antibody Treatment in Mice. Cancer Immunol Res, 2015. 3(6): p. 631–40.

30. Hashimoto, M., et al., CD8 T Cell Exhaustion in Chronic Infection and Cancer: Opportunities for Interventions. Annu Rev Med, 2018. 69: p. 301–318.

31. Lugli, E., et al., Stem, Effector, and Hybrid States of Memory CD8(+) T Cells. Trends Immunol, 2020. 41(1): p. 17–28.

32. Mootha, V.K., et al., PGC-1alpha-responsive genes involved in oxidative phosphorylation are coordinately downregulated in human diabetes. Nat Genet, 2003. 34(3): p. 267–73.

33. Bertrand, F., et al., TNFalpha blockade overcomes resistance to anti-PD-1 in experimental melanoma. Nat Commun, 2017. 8(1): p. 2256.

34. Perez-Ruiz, E., et al., Prophylactic TNF blockade uncouples efficacy and toxicity in dual CTLA-4 and PD-1 immunotherapy. Nature, 2019. 569(7756): p. 428–432.

35. Higuchi, M. and B. Aggarwal, TNF induces internalization of the p60 receptor and shedding of the p80 receptor. J Immunol, 1994. 152(7): p. 3550–8.

36. Chin, Y.R. and M.S. Horwitz, Mechanism for removal of tumor necrosis factor receptor 1 from the cell surface by the adenovirus RIDalpha/beta complex. J Virol, 2005. 79(21): p. 13606–17.

37. Britton, G.J., et al., Defined microbiota transplant restores Th17/RORγt+ regulatory T cell balance in mice colonized with inflammatory bowel disease microbiotas. BiorXiv, 2019.

38. Kahn, O. and J.R. Giles, TOX transcriptionally and epigenetically programs CD8+ T cell exhaustion. Nature, 2019. 571: p. 211–218.

39. Im, S.J., et al., Defining CD8+ T cells that provide the proliferative burst after PD-1 therapy. Nature, 2016. 537(7620): p. 417–421.

40. Kamphorst, A.O., et al., Proliferation of PD-1+ CD8 T cells in peripheral blood after PD-1-targeted therapy in lung cancer patients. Proc Natl Acad Sci U S A, 2017. 114(19): p. 4993–4998.

41. Belkaid, Y. and O.J. Harrison, Homeostatic Immunity and the Microbiota. Immunity, 2017. 46(4): p. 562–576.

42. Aukrust, P., et al., Serum levels of tumor necrosis factor-alpha (TNF alpha) and soluble TNF receptors in human immunodeficiency virus type 1 infection--correlations to clinical, immunologic, and virologic parameters. J Infect Dis, 1994. 169(2): p. 420–4.

43. Ahn, E., et al., Role of PD-1 during effector CD8 T cell differentiation. Proc Natl Acad Sci U S A, 2018. 115(18): p. 4749–4754.

44. Belnoue, Protracted course of lymphocytic choriomeningitis virus WE infection in early life: induction but limited expansion of CD8+ effector T cells and absence of memory CD8+ T cells. J Virol, 2007.

45. Beyer, M., et al., Tumor-necrosis factor impairs CD4(+) T cell-mediated immunological control in chronic viral infection. Nat Immunol, 2016. 17(5): p. 593–603.

46. Bryl, E., et al., Down-regulation of CD28 expression by TNF-alpha. J Immunol, 2001. 167(6): p. 3231–8.

47. Faith, J., Identifying gut microbe-host phenotype relationships using combinatorial communities in gnotobiotic mice. Science Translational Medicine, 2014.

48. Hermans, I.F., et al., The VITAL assay: a versatile fluorometric technique for assessing CTL- and NKT-mediated cytotoxicity against multiple targets in vitro and in vivo. J Immunol Methods, 2004. 285(1): p. 25–40.

49. Contijoch, E.J., Gut microbiota density influences host physiology and is shaped by host and microbial factors. eLife, 2019.

50. McNulty, Effects of Diet on Resource Utilization by a Model Human Gut Microbiota Containing Bacteroides cellulosilyticus WH2, a Symbiont with an Extensive Glycobiome. PLOS Biology, 2013.

